# A sound strategy for homology modeling-based affinity maturation of a HIF-1α single-domain intrabody

**DOI:** 10.1101/2020.05.14.096149

**Authors:** Min Hu, Guangbo Kang, Xin Cheng, Jiewen Wang, Ruowei Li, Zixuan Bai, Dong Yang, He Huang

## Abstract

A sound strategy for computer-aided binding affinity prediction was developed for *in silico* nanobody affinity maturation. Venn-intersection of multi-algorithm screening (VIMAS), an iterative computer-assisted nanobody affinity maturation virtual screening procedure, was designed. Homology modeling and protein docking methods were used to substitute the need for solution of a complex crystal structure, which is expanding the application of this platform. As a test case, an anti-HIF-1α nanobody, VHH212, was screened *via* a native ribosome display library with a 26.6 nM of K_D_ value was used as the parent. A mutant with a 17.5-fold enhancement in binding affinity (1.52 nM) was obtained by using the VIMAS strategy. Furthermore, the protein-protein interaction of interface residues, which is important for binding affinity, was analyzed in-depth. Targeting HIF-1α can sensitize PDAC tumors to gemcitabine, which is a potential co-treatment method for pancreatic cancer patients. Under combined treatment, the cytotoxicity of gemcitabine on pancreatic cancer cell lines increased with the enhanced-affinity of an intrabody. Thus, this study provides a platform for universal, efficient and convenient *in silico* affinity maturation of nanobodies.

## Introduction

Caplacizumab is the world’s first Nanobody^®^-based medicine and has been launched in Europe and the U.S. for the treatment of acquired Thrombotic Thrombocytopenic Purpura (aTTP) (1). This breakthrough has increased the interest in developing single-domain antibody-based biotherapeutics (2). Nanobodies (Nbs) have been identified as candidates for replacing conventional antibodies due to their unique features with a smaller size, better thermostability, low immunogenicity, and high solubility (3–5).

Recombinant nanobodies are derived from the variable regions of the heavy chain of immunoglobulin in Camelids (6). They consist of four conserved frame regions (FRs) and three complementarity determining regions (CDRs) that are responsible for exclusively recognizing antigen, especially CDR3 (7). Generally, library-based display technologies, such as phage, ribosome, and yeast display, are ideal tools for the selection of antibody fragments (8, 9). Once a set of antibodies has been isolated, their binding affinity often requires further optimization for potential therapeutic application. Display methods are used to improve affinity of antibodies *via* affinity maturation *in vitro.* Nevertheless, they are laborious and time-consuming (10–12). In these cases, developing a technological platform for high-efficiency affinity maturation of nanobodies is an urgent problem that must be solved.

With the advent of computational experimental data acquisition and analysis, algorithm optimization, and computing power, several strategies of antibody affinity maturation *in silico* have been developed based on structure-based rational design (13–17). Remarkably, E.O. Purisima’s group developed a platform and employed this for a nanobody, which resulted in an equilibrium dissociation constant (K_D_) of 2 nM; this led to a 9.4-fold improvement in binding affinities (17). The key feature of this platform was that high-affinity mutants could be obtained cost-and time-effectively.

In our previous study, the three-dimensional structure of a nanobody was constructed by homology modeling, while complex structures were gained by molecular docking (18). The outcome of such studies also helped us to gain better insight into antibody-antigen interactions and structures. In general, structure-based rational design is based on molecular mechanics with the addition of implicit solvent models, such as molecular mechanics-Poisson-Boltzmann surface area (MM-PBSA), generalized Born surface area (MM-GBSA) (19) or the Lazaridis-Karplus solvent model (MM-LKSM). On this basis, effective computer-assisted affinity maturation can be realized by advances in computational performance and force-field parameterization. Based on an assessment of binding free energy by different algorithms, a novel virtual screening strategy VIMAS was developed in the present study.

In this study, we explored the applicability of rational mutation hotspot design protocol (RMHDP) and Venn-intersection of multi-algorithm screening (VIMAS) strategies for nanobody affinity maturation, while continuing our efforts towards improved therapies of gemcitabine-resistance in PDAC. Although pancreatic cancer represents 3% of all cancers, the 5-year survival rate of PDAC patients is less than 9% (20). Recently, the PAXG arm has extended the survival of pancreatic cancer patients, but drug resistance to gemcitabine chemotherapy in pancreatic cancer has yielded unsatisfactory results (21). Clinical trials and pre-clinical studies indicate that targeting HIF-1α is an efficient strategy to reduce gemcitabine resistance (22). Chemical HIF-1α inhibitors lead to side effects and microenvironment modulation effects, which has restricted their application in clinical trials (23). Moreover, HIF-1α is an intracellular target; the applications of the full-length antibody are limited (24). In this case, nanobody-based intrabodies could provide a highly specific strategy for combined therapy.

## Results

### Construction of ribosome display library

A ribosome display library was constructed with a size of approximately 3.01×10^14^ individuals. Total RNA was extracted from lymphocytes of peripheral blood (see Supplementary Figure S1A), and a cDNA library was constructed by reverse transcription PCR. The peptide gene of interest was integrated with a T7 promoter, a stem-loop, and a translation initiation sequence at its 5’ site by nested PCR (see Supplementary Figure S1B, C). At the same time, a spacer sequence (gene III of the filamentous phage M 13, see Supplementary Figure S1D) and a stem-loop were added in its 3’ site. The ribosome display library was constructed with a linking peptide gene and gene III by overlap-PCR (see Supplementary Figure S1E). Colony PCR (see Supplementary Figure S1F) results showed that the target band was approximately 450 bp.

### Selection of HIF-1α-specific VHH

After four consecutive rounds of panning, 215 clones were isolated to evaluate the binding affinity by phage ELISA; BSA was chosen as negative control. Positive clone sequences were analyzed, which showed that they were identical and were able to be translated correctly (Figure 1). In conclusion, five different nanobodies (VHH67, VHH98, VHH167, VHH189, and VHH212) were identified from the ribosome display library. Based on this condition, VHH212 was selected for further optimization and *in vitro* affinity maturation (Figure 2A).

**Figure 1.**
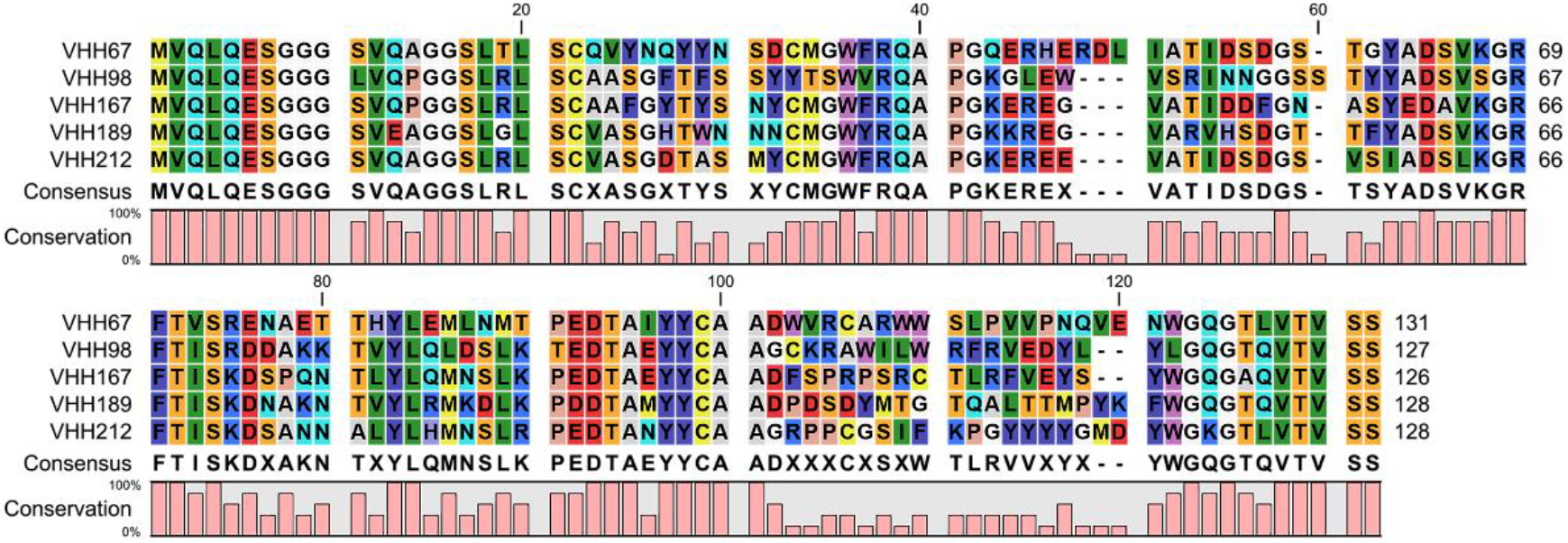
Alignment of the amino acid sequences of the positive clones of the nanobodies screened by phage ELISA.

**Figure 2.**
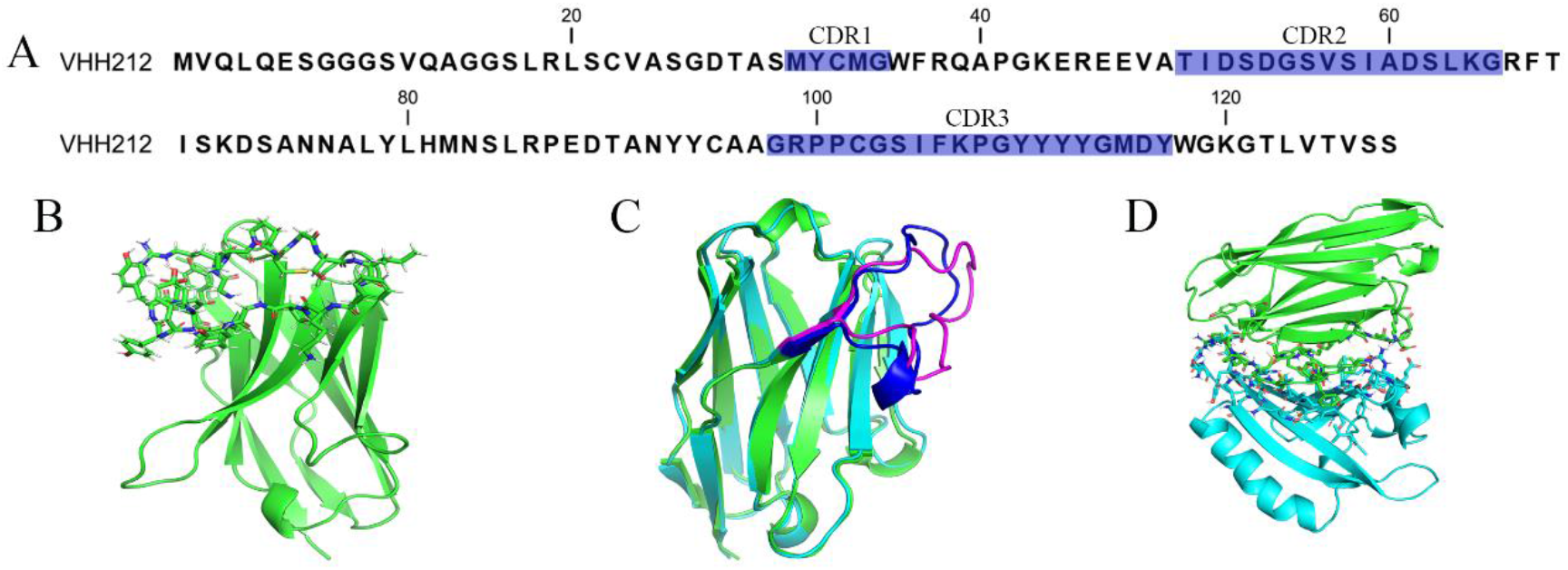
Amino acid sequence and three-dimensional structure of VHH212-07. (A) The framework and complementarity-determining region (CDR) sequences are defined according to the Kabat numbering scheme using the AbNum. (B) The three-dimensional crystal structure of VHH212-07 constructed by MODELLER. (C) VHH212-07 conformational alignment before (green with blue CDR3) and after (cyan with magenta CDR3) simulation. (D) Docked protein complex of HIF-1α-PAS-B (crystal structure in cyan) and anti-HIF-1α-PAS-B antibody VHH212-07 (homology model in green). Interfacial residues (5.0 Å) are shown in sticks representation.

### Homology modeling and MD simulation

Initially, the sequence alignment results are shown in figure S2B. The predicted models were validated based on their Verify 3D values, ERRAT plot and Ramachandran plot using the online server SAVES *v*5.0. Model 7 (marked VHH212-07) scored excellent results in all of the assessments. The Verify3D values for VHH212-07 show that at least 80% of the amino acids scored >= 0.2 in the 3D/1D profile. The Ramachandran plot of main residues showed that all residues were located in the most favored regions or additional allowed regions by 93.3% and 6.7% respectively (Figure S3A). Also, the Gly and Pro residues were located in allowed regions (see Supplementary Figure S3B, C). The three-dimensional structure of VHH212-07 is shown in figure 2B.

An MD simulation of 50 ns for the VHH212-07 structure was performed to refine the predicted structure. As shown in figure 2C, significant conformational rearrangements occurred in CDRs, especially in CDR3. The convergent energy curve shows that the potential energy had distinctly reduced (see Supplementary Figure S3D). Based on the analysis of backbone root-mean-square deviation of trajectory (Figure S3E), the structure shows minimal RMS fluctuations after 20 ns. This indicates that the structure reached equilibration and that the resultant trajectory is a stable conformation. Also, the root-mean-square fluctuations revealed small fluctuations of the residues that correspond to the loop region (see Supplementary Figure S3F). Thus, a high-quality homology model with stable conformation was obtained by MD simulation, which was used for molecular docking to identify possible active site residues.

### Docking and interfaces residues analysis

Based on the crystal structure of antigen (PDB ID: 4H6J) (25), antigen-antibody complexes were obtained by docking with HADDOCK. Ten clusters were provided with a low Z-score, which indicates how many standard deviations from the average this cluster is located in terms of score (see Supplementary Figure S3G). The complex structure with the lowest score and binding free energy was selected and analyzed visually using PyMol (Figure 2D).

For affinity maturation, InterProSurf was used to identify interface amino acid residues of the complex. Protein-protein interactions of the complex were analyzed, and a list of interface amino acid residues was predicted (see Supplementary Table S2) based on ASA and propensities most likely to be responsible for protein interaction. At the same time, the interface area and a change in the surface area of each residue upon complex formation were also provided. Considering that one method alone may lead to the omission of critical binding residues, we counted interface amino acid residues of VHH212 within 5Å of HIF-1α-PAS-B by PyMol (see Supplementary Table S2).

### *In vitro* affinity maturation

To maximize the chance of identifying mutations that increase antibody-antigen binding affinity, a VIMAS strategy was established by the combination of three separate computer algorithms for mutagenesis: mCSM-AB, OSPREY, FoldX. Initially, eleven sites were narrowed down as described in the methods. After that, the VIMAS strategy was employed to screen mutants with improved binding affinity in the first round, which greatly reduced the workload for next round screening.

With the use of former research results, a few numbers of residues on the protein-protein interface were selected as interesting targets for affinity maturation *in vitro,* which typically contribute to the stabilization of a complex (26, 27). In this case, eleven amino acid sites (four in CDR2 and seven in CDR3) that may alter antibody-antigen binding affinity were chosen by RMHDP. In the first round, 187 single-point mutations were evaluated by three protocols (mCSM-AB, OSPREY and FoldX) (Figure 3A). The calculation results of the first round are shown in table S3, S4 and S5, respectively. Eventually, thirteen single-point mutations in three amino acid positions (A60, I105, K107) were narrowed down by this process for further evaluation (Figure 3B). Based on the positive results of single-point mutations, double and triple mutants were introduced to the next round of calculations.

**Figure 3.**
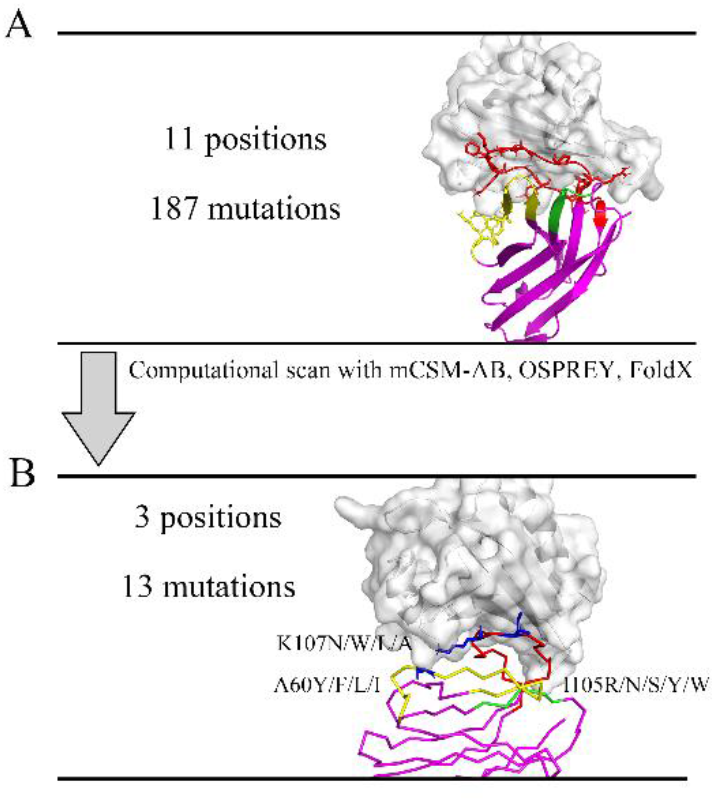
First-round screening leading to validated affinity-improved single-point mutants. (A) The proximity of virtually scanned CDR residues to the HIF-1α-PAS-B epitope. (B) Three-dimensional locations of single-point mutations with binding affinity improvement. It shows the mutation sites we selected Ala60, Ile105 and Lys107. The antigen is shown as gray ribbon in a translucent molecular surface. The VHH is rendered as a magenta ribbon/Cα-trace, with CDR1 in green, CDR2 in yellow, and CDR3 in red. Models of mutated side-chains are shown as blue sticks.

For the second round, 56 double mutants were formed by exhaustively combining thirteen single-point mutations and were verified computationally within FoldX. Our calculation shows that most double-point mutants showed more negative binding free energy than single-point mutants (see Supplementary Table S6). This indicates that the affinity increases caused by single-point mutations have an additive effect. For example, when two sites undergo amino acid mutations such as Ala60 to Phe and Lys107 to Leu, the difference is optimal for decreasing the binding free energy (−2.28 kcal/mol); thus, the mutated antibody is more stable than the parental antibody. However, the binding free energies were only −0.82 kcal/mol and −0.99 kcal/mol, respectively, when A60F and K107L mutations were carried out independently (Figure 4).

**Figure 4.**
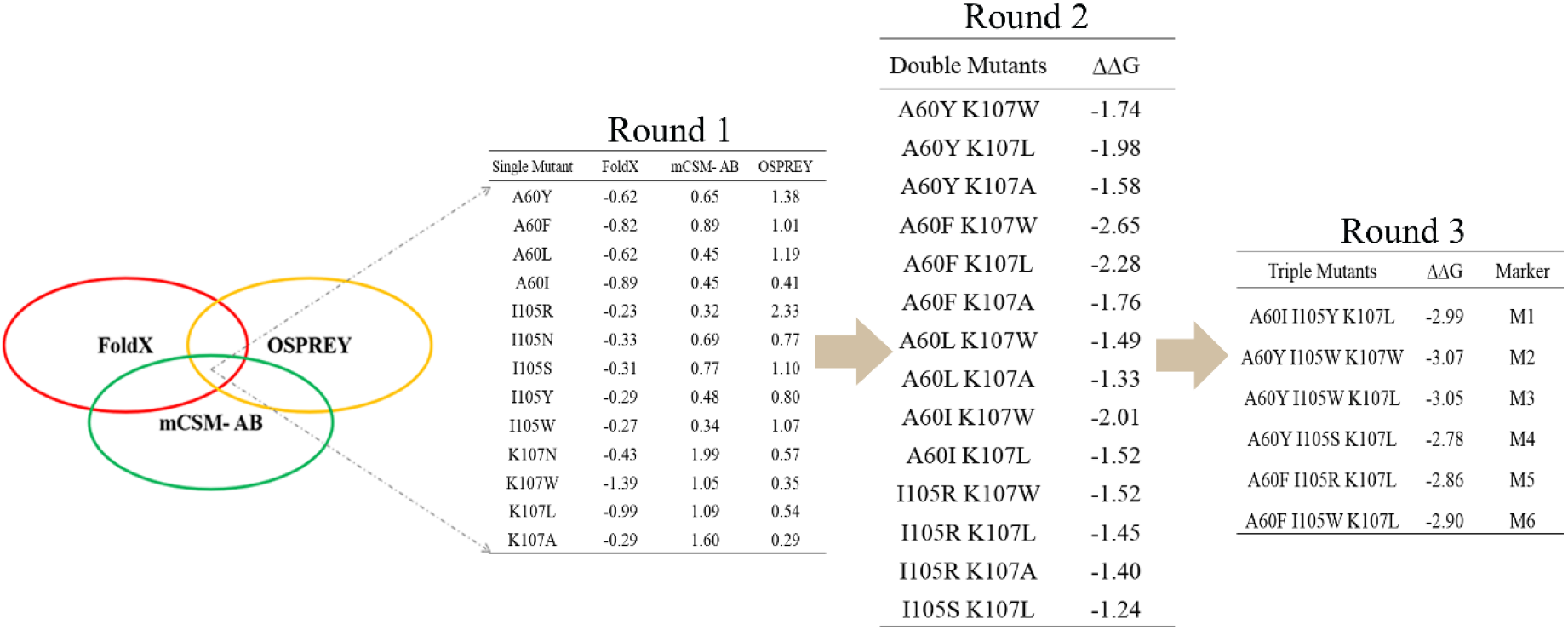
Flow chart of the VIMAS strategy. Totally three rounds iterative computer-assisted virtual screening was performed, while six triple mutants with enhanced-affinity were obtained. The round 1 shows the results of intersection in the Venn diagram. They are thirteen single point mutants screened by three scoring functions. The thirteen single point mutants are not ordered, and they’re chosen because they stand out in all three scoring functions. As shown in the round 1, the second column is the binding free energy change (ΔΔG) after mutation by FoldX; the third column is antibody-antigen affinity change upon mutation by mCSM-AB; the fourth column is the change of K* Score (log10) calculation by OSPREY.

For the third round, 55 triple mutants were formed by exhaustively combining fourteen double-point mutations. Most triple mutants showed better binding affinity than double mutants (see Supplementary Table S7). Ultimately, six triple mutants were selected and named M1-M6 (Figure 4).

### Binding measurements to identify affinity-matured mutants

As described in methods, the parent and six predicted mutants were synthesized, expressed and purified. As shown in figure 5A, purified VHHs presented a clear single band at approximately 15 kDa on a 15% SDS-PAGE gel. ELISA results revealed that all mutants showed a stronger binding signal than the parent. Comparing with parent, the binding affinity of mutants M3 and M1 significantly increased (Figure 5B). BSA was employed as a negative control, which showed no change on response value as the concentration increased.

**Figure 5.**
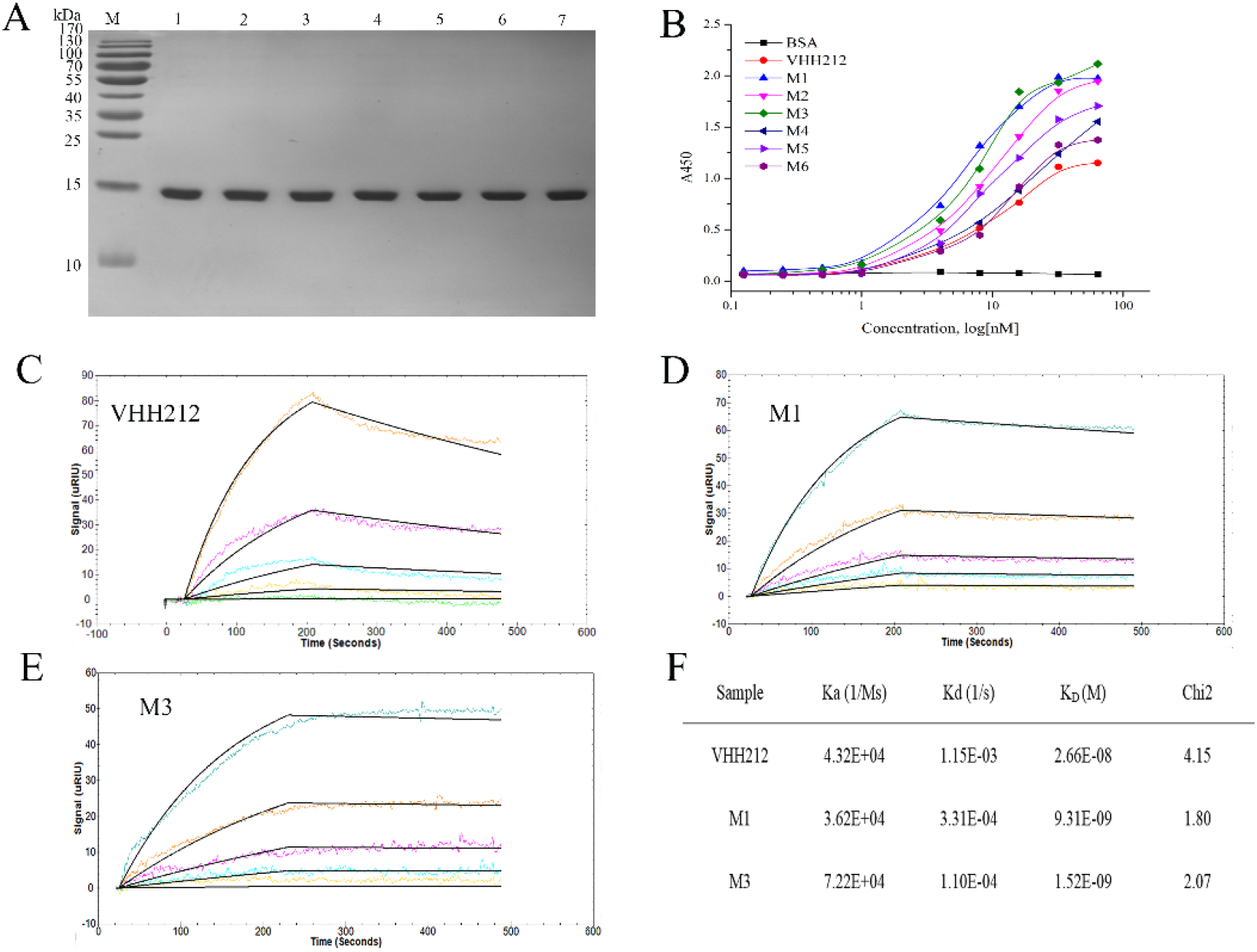
Analysis of binding affinity and thermostability of VHHs. (A) SDS-PAGE analysis for purified VHHs. M, molecular weight marker; 1, VHH212; 2-7, M1, M2, M3, M4, M5, M6. All samples were loaded to 7 μg. (B) ELISA assay was used to reveal the binding affinity improvement of the mutants. All mutants and parental nanobody were serially diluted and tested for binding affinity to wells coated with antigen. The BSA was used as negative control. (C) Representative SPR sensorgram binding profiles. Interaction of the parent VHH212, the lead triple mutants M1 (D), and the lead triple mutant M3 (E) with immobilized antigen. The color lines represent raw data, and the black lines are global fits to a 1:1 bimolecular interaction model. (F) SPR K_D_ values of parental VHH212 and mutants.

To further demonstrate that the predicted mutants indeed conferred increased binding affinity, K_D_ values were determined by 1:1 surface plasmon resonance measurements using a Reichert 4SPR instrument. Figure 5C, D and E show the sensorgrams for parental VHH212 and mutants. Compared with parental VHH212, mutants showed higher binding affinities. By contrast, M3 showed a 17.5-fold increase in binding affinity, which arose from a small 1.67-fold increase in Ka coupled with 10.5-fold hugely decrease in Kd compared with the parent nanobody. All the values of SPR sensorgrams are shown in figure 5F.

### IC_50_

As described, VHH212, M1 and M3 were transfected into human pancreatic cancer cell lines MIA PaCa-2, and BxPC-3, respectively. An IC_50_ assay was performed to evaluate the effect of combination treatment. As shown in figure 6A and B, compared with the vector control, targeting HIF-1α therapy increased the cytotoxicity of gemcitabine. Addition of VHH212, M1 or M3 intrabody was associated with a decrease in the gemcitabine IC_50_ dose from 62.73 to 41.43, 32.49 and 32.8 nM for BxPC-3, respectively. Intrabody was associated with a decrease in the gemcitabine IC_50_ dose from 59.73 to 41.78, 22.9 and 22.66 nM for MIA PaCa-2, respectively. In summary, these data suggest that optimized mutants could translate into a significant enhancement of HIF-1α neutralization at the cellular level.

**Figure 6.**
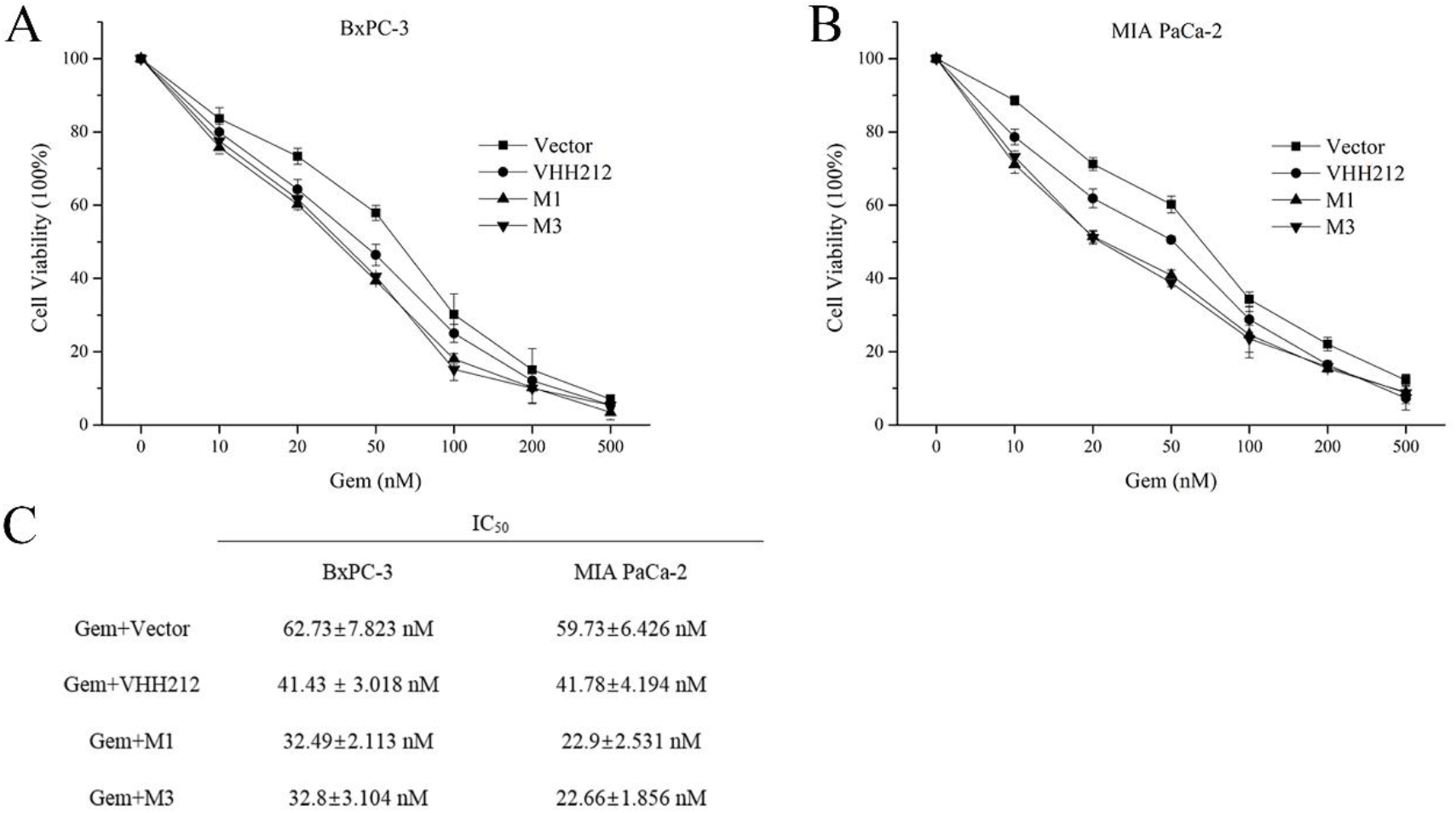
Nanobody-based intrabody could enhance the cytotoxicity effects of gemcitabine on pancreatic cancer cell lines. *In vitro* cytotoxicity assay show that nanobody-based intrabody could enhance the antitumor and cytotoxicity effects of gemcitabine on pancreatic cancer cell lines, BxPC-3 (A) and MIA PaCa-2 (B), respectively. (C) IC_50_ values were calculated using GraphPad Prism 7.0 software. Data are the mean ± SD of triplicate determinations.

### Structural insight of the affinity-improving mutations

As shown in figure 7, the in-depth analysis of the three-dimensional structure of the M3 and HIF-1α-PAS-B domain complex helps understand the theory of the protein-protein interaction of interface residues, which is valuable for further studies of increasing binding affinities. The antibody-antigen complex was analyzed before and after maturation. A salt bridge between D61 and R258 with a distance of 3.85 Å was present in the original complex, while the distance became 3.28 Å after Ala was substituted with Tyr at position 60 (Figure 7A). Furthermore, a salt bridge was formed between E47 and R258. At the A60 site, mutation to Y60 generates a hydrophobic interaction with L243 and Y254 of antigen (Figure 7B). We also found that a hydrophobic effect was formed at sites 105 and 107 after mutation (Figure 7C). The add in of new hydrophobic interactions at these three positions was likely responsible for protein structural stabilization. The mutation of A60Y generates hydrogen bonds: the main chain-side chain H-bound between A60Y and E257 with a distance of 2.5 Å (Figure 7D). Figure 7E shows the cation-Pi interaction between A60Y and R258. Additionally, the mutation of A60 to Y introduces new aromatic-aromatic interactions with Y254 of HIF-1α (Figure 7E). The substitution of I with W at position 105 also formed an aromatic-aromatic interaction with Y340. Additionally, mutations at positions 105 and 107 led to a smaller distance between F106 and Y340 with 3.91 Å of aromatic-aromatic interactions (Figure 7F). The introduced of aromatic residues generated aromatic-aromatic interactions that played a pivotal role in binding affinity improvement.

**Figure 7.**
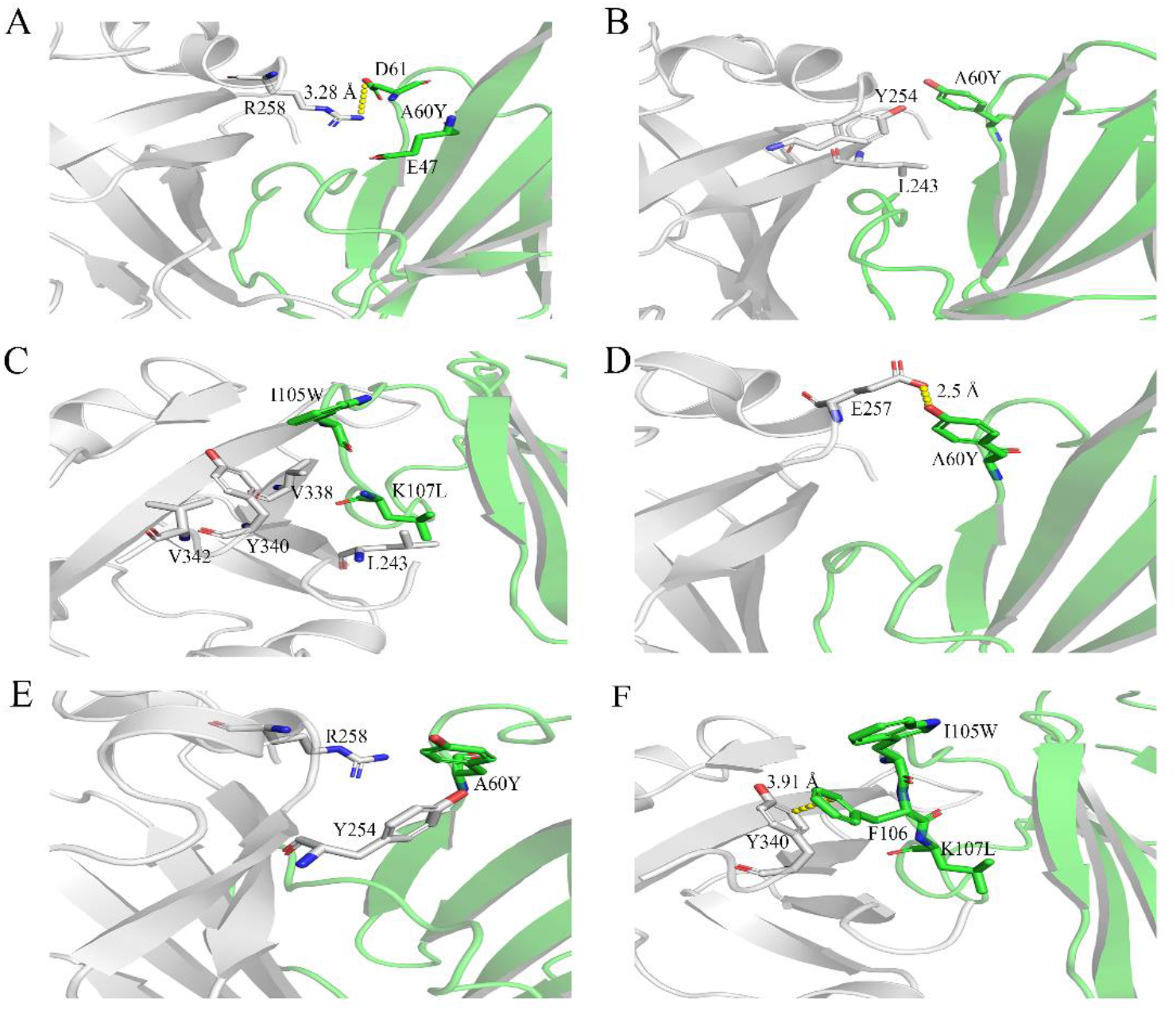
Structure models of optimized antigen-M3 interactions. The M3 and HIF-1α-PAS-B chains are colored green and grey, respectively. The corresponding N atoms are colored blue and O atoms colored red. (A) A salt bridge is formed between D61 and R258 of HIF-1α-PAS-B. (B) Structure model of the A60Y. The new hydrophobic interactions are created between A60Y and L243 of HIF-1α-PAS-B. (C) Hydrophobic interactions formed by I105W and K107L. (D) A60Y introduced new Hydrogen-bond with E257. (E) Cation-Pi interaction (A60Y-R258) and Aromatic-Aromatic interaction (A60Y-E254). (F) Aromatic-Aromatic interaction of I105W and Y340.

## Discussion

With the first nanobody-based medicine Caplacizumab launched in Germany, the applications of nanobodies have stimulated research interest towards the development of nanobody-based biological drugs. Based on the characteristics of VHHs, nanobodies should not cause ADCC. High binding affinities are also essential for nanobody-based biotherapeutics to compete with the natural sub-units or receptors. Nevertheless, the binding affinity of nanobodies screened by display technology tends to be weaker than those derived from mammalian platforms, which limits their application in drug development (28, 29). In this case, developing a method or a sound strategy to focus on nanobody affinity maturation is an urgent problem that must be solved. With the growth of antibody-antigen complex structural databases and the development of algorithms for binding affinity prediction, computer-aided rational design has been a tractable choice for the maturation of antibody affinity characteristics (30–33).

Here, we designed a sound strategy for universal, efficient and convenient computer-aided rational design, which was used to significantly increase the binding affinity of a nanobody. Many successful computer-aided affinity maturation cases have been reported in the early stage, which is a highly effective strategy that can often lead from 10 to 100-fold increases in binding affinity (16, 18, 34). Herein, a three-dimensional structure developed by homology modeling and docking was used instead of crystallographic structures, which broadens application scenarios for our strategy. RMHDP provided an effective strategy to select binding sites with a significant effect on affinity. This was useful for predicting interface residues and reducing the number of calculations. VIMAS is a protocol designed explicitly for nanobody affinity maturation *in vitro*. Based on the low molecular weight of nanobodies, and considering the practicability of using three different algorithms, this approach was chosen for our calculations. These algorithms are being made open-source to provide a convenient, user-friendly platform for other researchers.

Hypoxia is a typical characteristic in the microenvironment of solid tumors that is linked to drug resistance in various cancers (35, 36). Previous studies have indicated that HIF-1α is responsible for tumor cells’ resistance to cisplatin, oxaliplatin, gemcitabine and paclitaxel (37, 38). In this case, targeting HIF-1α was perceived as an efficient strategy for treating pancreatic cancer (22, 39). Our study indicated that anti-HIF-1α VHH212 has anti-tumor effects by sensitizing cells to gemcitabine cytotoxicity *in vitro*. Moreover, with the enhancement of the binding affinity of the nanobody, the capacity for neutralizing antigen epitopes and competitive inhibition of the formation of dimers increased simultaneously. Remarkably, we provided a potential co-treatment method for solving the issue of drug resistance to gemcitabine chemotherapy in pancreatic cancer patients.

In conclusion, we develop a sound strategy for computer-aided nanobody affinity maturation. The protein-protein interaction of interface residues was analyzed in-depth. The feasibility of the strategy was validated by the acquisition of a mutant with a 17.5-fold increase in binding affinity (1.52 nM) is obtained. Importantly, the nanobody-based intrabody could enhance the antitumor and cytotoxicity effects of gemcitabine in pancreatic cancer cell lines. The shortcomings of the *in silico* affinity maturation research are currently limited by computing power, structural databases, and algorithms for binding affinity prediction. For a long time, the development of a virtual screening strategy based only on the three-dimensional structures of antigens has been the most challenging and important goal of research in this exciting field.

## Experimental Procedures

### Construction of ribosome display library

A naïve library was constructed by amplifying and cloning the Nanobody™ repertoire according to the method described by Christian Zahnd (40). Four peripheral blood samples from two Bactrian camel were collected as the source of lymphocytes using a lymphocyte isolation kit (Thermo Fisher Scientific). Total RNA was extracted, and the cDNA was synthesized with the total RNA as template and oligo (dT) 18 as a primer using a First Strand cDNA Synthesis Kit (Thermo Fisher Scientific). Next, the cDNA was amplified in three steps by PCR. The primers used in all the PCRs are shown in table S1. Finally, the PCR products were identified by gel electrophoresis, digested with *NcoI* and *NotI* restriction enzymes, and ligated into a pCANTAB 5E vector (Biofeng). *Escherichia coli* TG1 cells were electroporated (2.5 kV, 25 μF, 200 Ω) with the linked products using an Eppendorf Eporator^®^ instrument (Eppendorf). Colony PCR and sequence analysis (GENEWIZ) were performed to identify library diversity.

### Selection of HIF-1α-specific VHH

For the selection of nanobodies against HIF-1α, a Nunc Maxisorp^®^ 96-well plate was coated with recombinant HIF-1α-PAS-B at 0.1 mg/mL in a coating buffer overnight at 4 °C and was washed three times with PBSM (PBS buffer with 5 mM MgCl2), then blocked with a blocking buffer for 2 h at room temperature. Wells were then washed, and the completed translation library was added to the plate wells. Further, the plate was washed three times, and the mRNA of bound was eluted with 200 μL of elution buffer with incubation in a water bath at 42 °C for 5 min. Finally, eluted mRNA was isolated and reversed transcribed to cDNA. After four rounds of screening, VHH gene fragments were inserted into a pCANTAB 5E vector and were transformed into *E. coli* TG1. Two hundred and fifteen individual clones were randomly chosen and detected by phage ELISA against HIF-1α-PAS-B and BSA. The positive VHH clones responded by absorbance at OD450 were selected and sequenced by GENEWIZ.

### Homology modeling

Based on the previous screening, we obtained a nanobody VHH212 (accession No: MK978327) with high binding affinity. To find suitable templates for modeling, we submitted our sequence to Protein BLAST (41) for searching. A total of five templates, namely 3STB (42), 4DK3 (43), 4W6Y (44), 5IMK (45), and 5BOP (46), were selected based on high sequence identity (>65%). Furthermore, the query coverage and E-values all meet the requirement for templates. Three-dimensional structures of the parent VHH212 were developed as described in ref. (18). The predicted models were assessed using web server SAVES *v*5.0 (http://services.mbi.ucla.edu/SAVES/), which contained Ramachandran plots, Verify3D and ERRAT plots.

### Molecular dynamics simulation

The VHH212 model was performed molecular dynamics (MD) simulation in GROMACS *v*4.5.5 (47), and an AMBER99SB-ILDN force field was applied to generate the topology file. The model was located in a dodecahedron box at a distance of 1.2 nm from the box’s edges, and solvated with SPC water. Na^+^ and Cl^-^ (0.15 M) were added to neutralize the systems. Energy minimization was carried out to relax the internal constraints. After minimization, the system was equilibrated by NVT and NPT for 100 ps. Through the modified Berendsen thermostat, the system temperature was controlled at 300K with a coupling time of 0.5 ps (48). Meanwhile, the pressure was maintained at 1 bar with coupling constants of 1.0 ps by Parrinello-Rahman isotropic pressure coupling (49). The time step for integration was 2 fs with LINCS constraint algorithm. Periodic boundary conditions were used, and a production run was carried out for 50 ns of simulation. Finally, the analysis was implemented in the Gromacs package.

### Docking and RMHDP

The three-dimensional structure of HIF-1α-PAS-B was obtained from the PDB database (PDB ID: 4H6J). Initially, antibody and antigen models were prepared for using with HADDOCK. All water molecules were removed, and H-atoms were added to the protein crystal structure using PyMol (http://www.pymol.org/). VHH212 was docked to antigen with the web “easy interface” of HADDOCK (50), default parameters were used. The CDR of the antibody was defined as active residues. The active residues of antigen were determined based on related studies (51–53). Eventually, the residues 242-262, 299, 304, 306, 308, 318-327 and 333-342 were chosen. The residues of antibody and antigen adjacent to the active site were defined as passive sites.

RMHDP strategy was employed to effectively screen interface residues (18). At the same time, InterProSurf (54) was used to predict interacting residues of a protein complex. The complex model file was uploaded to the server. InterProSurf printed out interface residues, interface area and the surface area change of each residue as it forms a complex by analyzing each chain within the complex. Furthermore, the residues of the protein within 5 Å of the ligand were counted using PyMol.

### *In silico* affinity maturation

VIMAS, a new strategy specific for nanobody affinity maturation *in silico,* was proposed. Three protocols (mCSM-AB, OSPREY, and FoldX) were used to predict the effect of mutations on affinity. Each amino acid was sequentially mutated to each of 18 amino acids (excluding Cys and Pro), and the binding affinity change relative to the parent nanobody was calculated. Similarly, the antibody-antigen complex was refined according to the requirements of each method.

#### mCSM-AB

mCSM-AB is a user-friendly web server freely available at (55). It relies on graphbased signatures to predict antigen-antibody affinity changes upon mutation while considering the dataset of the AB-Bind Database (30) in order to assess the applicability of the signature. The antibody-antigen complex was uploaded, and the mutations of the antibody were informed. The mutation information needs to be given, including residue position, wild-type and mutant residue codes in one-letter format and chain identifier. The amino acid residues of the antibody interface interacting with antigen analyzed above were mutated to others.

#### OSPREY

OSPREY *v* 3.0 beta, a protein design software with new python interface, was employed to predict the impact of protein mutation on ligand binding (56). Typically, the initial input complex structure must be modified to make it compatible with OSPREY. Firstly, the residues of missing atoms can be deleted from the structure if it does not affect mutations. Otherwise, model the missing atoms in a reasonable conformation. Secondly, make sure hydrogens had been added in the input structure. Finally, all His protonation states must be considered; OSPREY recognizes three different His residues states. When the hydrogens had been provided in the input structure, the auto-fix feature will rename histidine residues by MolProbity server (57). The residues were systematically mutated to all others as the description of the Python script with the BBK* design in the OSPREY package.

#### FoldX

The <BuildModel> module of FoldX was used to estimate the stability effects of single-point mutations (58). It can be run in YASARA to analyze protein-protein interaction energies and calculate the free energy differences between the designed mutants and parental antibodies. Each mutation was calculated three times following the recommended protocol (pH 7, temperature 298 K, ion strength 0.050 M, VdWDesign 2). The changes of antibody-antigen complex in binding free energy (G, kcal/mol) was calculated. The optimal amino acid mutation was selected based on the extent to which its binding free energy was reduced.

### Protein expression and purification

The DNA sequence of parent VHH212 and mutants were synthesized and subcloned into a pET-32a (+) expression vector, with an N-terminal RBS/TATA box and C-terminal 6×His tag, by GENEWIZ. The vectors were then transformed into *TransB* (DE3) competent cells (TransGen) according to the manufacturer’s instructions and were selected on ampicillin (50 μg/mL) agar plates. Purification was performed by combing a HisTrapFF column and analytical size-exclusion chromatographic Superdex™ 75 5/150 with an ÄKTAexplorer 100 (GE Healthcare). The human cDNA sequence for the HIF-1α-PAS-B domain was synthesized and subcloned into the prokaryotic expression vector pGEX-4T-1 (Amersham). The recombinant protein was produced in Rosetta (DE3) cells (Novagen) and purified by affinity chromatography using a GSTrapFF column. The elution fraction was identified by 15% SDS-PAGE electrophoresis.

### ELISA

The HIF-1α-PAS-B domain was coated at 3 μg/ml onto 96-well microtiter plates (Thermo Fisher Scientific) and incubated overnight at 4 °C. After three washes with PBST (PBS buffer with 0.05% Tween 20), residual binding sites were blocked using 5% skimmed milk in PBS for 2 h at 37 °C. The blocking buffer was decanted, and antibodies were diluted in PBS then added to the wells. After five washes in PBST, a secondary antibody of HRP-conjugated anti-6×His (Abcam) was added at a 1:5000 dilution and incubated for 1 h at 37 °C. The binding was detected using a TMB Two-Component Substrate solution kit (Solarbio^^®^^) for 20 min at 37 °C, and the reaction was terminated by adding 1 M H2SO4. Absorbance signals were measured at a wavelength of 450 nm in Synergy HT (Bio Tek).

### SPR

All binding experiment assays were performed by Reichert 4SPR (Reichert Technologies). The recombinant HIF-1α-PAS-B was captured onto Planar Mixed SAM sensor chip (catalog no. 13206061) using high affinity binding of biotin to avidin until a signal increase of approximately 1000 micro-refractive index units (μRIUs) was achieved. The running buffer solutions used in all experiments was PBST. Nanobodies (3.125-200nM) were injected over the chip at a flow rate of 25 μL/min with 180 s contact time and 300 s dissociation time. The regeneration of chip was carried out with 10 mM (pH 2.5) Glycine-HCl with 90 s injection. The background response was recorded using the reference flow channel, and the background data was subtracted from the data from each injection sample. The kinetic data were analyzed using *TraceDrawer* software (Ridgeview Instruments) with a 1:1 binding model. All mutant and parental VHH212 samples were run in duplicates.

### Intrabody inhibition assays (IC_50_)

MIA PaCa-2 and BxPC-3 human pancreatic cancer cell lines were seeded into 96-well plates at a density of 3000 cells/well. After 24 hours incubation at 37 °C under hypoxic condition, 200 ng of pEGFP-N1, pEGFP-N1-VHH212, pEGFP-N1-M1 and pEGFP-N1-M3 plasmids were transfected into cancer cell lines using Lipofectamine^TM^ 3000 (Invitrogen). Six hours after transfection, fresh medium containing different concentrations of gemcitabine (0, 10, 30, 50, 100, 200, 500 nM) was added into a 96-well plate. After 48 hours of incubation, 10 μL CCK-8 (Solarbio) was added to a 96-well plate. After two additional hours of incubation at 37 °C with CKK-8, absorbance was measured at 450 nm by Synergy HT (Bio Tek). Half-maximal inhibitory concentration (IC_50_) was analyzed relative to the blank control using Graphpad 7.0. Values are shown as the means of triplicate wells from three independent experiments for each drug concentration.

### Data availability

All data are contained within the article and accompanying supporting information.

## Funding and acknowledgments

The present study was supported by grants from National Key Research and Development Project (No. 2019YFA0905600), National Natural Science Foundation of China (No. 31470967), Science and Technology Program of Tianjin, China (No. 19YFSLQY00110), Natural Science Foundation of Tianjin, China (No. 19JCTPJC40900). We would like to thank MogoEdit to minimise incorrect English expressions.

## Footnotes

The abbreviations used are: PDAC, pancreatic ductal adenocarcinoma; PAXG, cisplatin, nab-paclitaxel, capecitabine, gemcitabine combined treatment arm; ASA, accessible surface area; MD, molecule dynamics; NVT, Number of particles, Volume, Temperature; NPT, Number of particles, Pressure, Temperature; LINCS, Linear Constraint Solver; BSA, bovine serum albumin; SPR, surface plasmon resonance; ADCC, antibody-dependent cell-mediated cytotoxicity.

